# Instability in computational models of vascular smooth muscle cell contraction

**DOI:** 10.1101/2023.10.16.562505

**Authors:** Alessandro Giudici, Jason M Szafron, Abhay B Ramachandra, Bart Spronck

**Affiliations:** Department of Biomedical Engineering, CARIM School for Cardiovascular Diseases, Maastricht University, Maastricht, The Netherlands; GROW School for Oncology and Reproduction, Maastricht University, Maastricht, The Netherlands; Department of Pediatrics, Stanford University, Stanford, CA, United States; Department of Biomedical Engineering, Yale University, New Haven, CT, United States; Macquarie Medical School, Faculty of Medicine, Health and Human Sciences, Macquarie University, Sydney, NSW, Australia

**Author notes:** **Corresponding author:** Bart Spronck, PhD, Department of Biomedical Engineering, Cardiovascular Research Institute Maastricht (CARIM), Maastricht University, Universiteitssingel 40, Room C5.578A, Maastricht, 6229 ER, The Netherlands. Authors contributed equally (Alessandro Giudici, Jason M Szafron).

**Keywords:** vascular smooth muscle cells, constitutive modelling, active stress modelling, instability, arterial mechanics, vascular smooth muscle cell contraction

## Abstract

**Purpose:** Through their contractile and synthetic capacity, vascular smooth muscle cells play a key role in regulating the stiffness and resistance of the circulation. To model the contraction of blood vessels, an active stress component can be added to the (passive) Cauchy stress tensor. Different constitutive formulations have been proposed to describe this active stress component. Notably, however, the *ex vivo* measurement of the biomechanical behaviour of contacted blood vessels presents several experimental challenges, which complicate the acquisition of comprehensive data sets to inform complex active stress models. In this work, we examine formulations for use with limited experimental contraction data as well as those developed to capture more comprehensive data sets.

**Methods:** We prove analytically that a subset of these formulations exhibits unstable behaviours (i.e., a non-unique diameter solution for a given pressure) in certain parameter ranges, particularly when contractile deformations are large. Furthermore, using experimental literature data, we present two case studies where these active stress models are used to capture the contractile response of vascular smooth muscle cells in the presence of 1) limited and 2) extensive contraction data.

**Results:** Our work shows how limited contraction data complicates the selection of an appropriate active stress model for vascular applications, potentially resulting in unrealistic modelled behaviours.

**Conclusion:** As such, the data presented herein provide a useful reference for the selection of an active stress model which balances the trade-off between accuracy and the available biomechanical information.

## 1. Introduction

Vascular smooth muscle cells (VSMCs) play a crucial role in regulating stiffness and resistance of the circulation [1,2]. The relative arrangement, distribution, and tone of VSMCs varies for different vascular segments [3–6]. Elastic vessels such as the aorta contain elastic lamellae separating concentric layers of VSMCs, while more distal muscular arteries such as the femoral artery contain fewer lamellae and thicker layers of VSMCs [4,6]. These structural differences reflect functional differences between arterial segments. In peripheral arteries and arterioles where resistance to blood flow is high, VSMC contraction allows for the regulation of tissue perfusion according to specific needs. In central arteries, which exhibit lower levels of vascular tone than peripheral vessels [3] and where resistance is low, VSMC contraction shifts load-bearing between arterial wall constituents and layers [5,7,8]. As such, VSMC contraction allows for the active modulation of arterial stiffness at any given pressure level [9–12]. Preserving the physiological VSMC function is key for maintaining vascular homeostasis and its disruption is linked to the development of vascular disease [1,5,13,14]. Therefore, quantifying changes in vascular contractility is critical to understanding disease development and identifying potential interventions.

In *ex vivo* settings, vascular contractility can be assessed by measuring changes in the biomechanical behaviour of a vessel in response to the administration of a vasoconstrictor, such as phenylephrine [5]. Because VSMCs are embedded in a microstructurally complex, radially heterogeneous extracellular matrix, the effect of their contraction on the vessel mechanics also depends on the passive mechanical behaviour of the arterial wall [15]. Furthermore, VSMC contractile capacity, as well as the passive mechanical behaviour, strongly depends on the loading configuration [16,17]. Vascular contractility characterisation must, therefore, be conducted in pseudo-physiological experimental conditions. Recent biaxial experiments have allowed for examining the VSMCs behaviour in a physiological loading environment with both pressurisation and axial extension of an intact vessel to an approximate physiological *in vivo* state [5,16]. Constitutive modelling can then be used to integrate the measured mechanical data [4,5], with the goal of linking changes in clinically relevant metrics with the underlying microstructure.

Despite the developments of sophisticated experimental set-ups for the biomechanical phenotyping of blood vessels, several considerations are necessary when investigating vasoconstriction in *ex vivo* settings. First, VSMC contraction is much slower than that of the skeletal or cardiac muscle. This has practical implications for *ex vivo* investigations (e.g., ∼15 min is a typical time between the *ex vivo* administration of a vasoconstrictor and the reaching of a stable level of vasoconstriction). Second, VSMCs are an active arterial wall constituent, which, as such, continuously respond to their chemical and mechanical environment. Therefore, investigating their behaviour over wide ranges of deformations is not trivial, as their contraction is continuously modulated in response to altered mechanical stimuli. Third, VSMC survival in excised vessels is limited in time and condition of storage, which further complicates experimental practices. Because of these considerations, the amount of mechanical data on the active VSMC contraction acquired experimentally is generally limited, which complicates informing computational models of active VSMC contraction. This is particularly relevant as some VSMC contraction models show unstable inflation behaviours for active contraction within the physiological range (**Figure 1**). Fourth and finally, typically, contraction experiments often contain limited mechanical information. For example, a typical experiment consists of keeping the vessel at a fixed axial stretch and pressure, and then inducing contraction [5,18]. Such contraction shifts the relationship between pressure and diameter [5]; however, only one point on this relationship is effectively recorded. As one of the goals of collecting contractility data is to capture material behaviour of the vessel constitutively for simulation of clinically relevant metrics or to predict changes over time under pathological loading conditions, care must then be taken in choosing an appropriate VSMC material model for the loading conditions to be simulated. Notably, the number of model parameters that can be uniquely fit from a typical experiment is usually small and limited by the experimental data available.

**Fig. 1.**
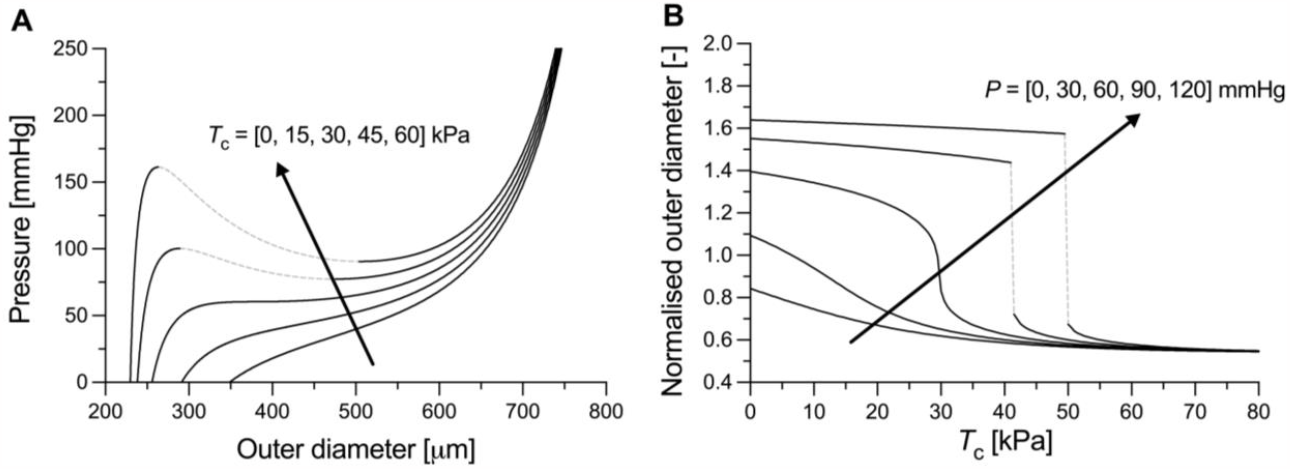
Simulated response to inflation at a constant (*in vivo*) value of axial stretch of the mouse descending thoracic aorta at growing levels of vascular smooth muscle tone. Instability regions are denoted by grey dashed lines. The passive behaviour of the mouse aorta (i.e., with circumferential active Cauchy stress 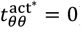) has been modelled using a four-fibre family strain energy density function according to data in Spronck et. (2021). The VSMC active contribution was modelled as a constant Cauchy stress 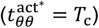 acting uniformly across the wall thickness. *P*: transmural pressure. Normalised outer diameter (Panel B) indicates the ratio between loaded and unloaded outer diameters. Direction of the arrow indicates increasing value of the parameter, i.e., *T*_c_ in panel A and *P* in panel B.

The aim of this work, therefore, is to examine different descriptive material models of active contraction available in the literature and understand their potential to yield appropriate deformation behaviour for a wide range of loading conditions and contractile deformations, evident in different regions of the vasculature. Models are investigated both analytically and numerically to understand their use in capturing experimental data.

## 2. Theoretical analysis (in a thick-walled cylinder)

In this section, we propose an analytical framework to evaluate the potential of active stress models to yield unstable contractile behaviours. While here this theoretical analysis is provided for a thick-walled arterial model, the Supplementary information illustrates how similar conclusions can be obtained when modelling arteries as a thin-walled cylinders.

For an axisymmetric, cylindrical tube, the only non-trivial linear momentum balance equation lies in the radial direction with

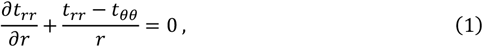

where **t** is the Cauchy stress tensor (with subscript *rr* and *θθ* indicating its radial and circumferential components) and *r* is the radial coordinate in the deformed configuration. This equation can be integrated to yield

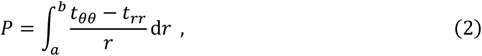

with transmural pressure *P* = *t*_*rr*_(*b*) − *t*_*rr*_(*a*), loaded inner radius *a*, and loaded outer radius *b* > *a*. Assuming an additive form for the active and passive stress contributions, the Cauchy stress tensor can be written as 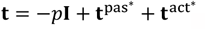. The passive part of the Cauchy extra stress 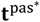 depends constitutively on the passive stored energy density Ψ^pas^ according to

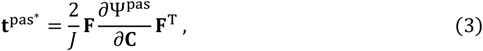

with deformation gradient tensor **F**, right Cauchy-Green deformation tensor **C** = **F**^T^**F**, and *J* = det **F** = 1 under the assumption of incompressibility as commonly assumed for vascular tissue. A Lagrange multiplier *p* is used to enforce the incompressibility constraint, which can be found generally for any position *r* within the wall as

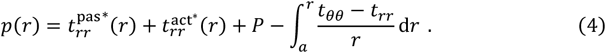

For pressurization and axial extension of a blood vessel in the absence of torsion, the deformation gradient tensor is **F** = diag[*λ*_*r*_, *λ*_*θ*_, *λ*_*z*_] with *λ*_*θ*_ = *r*/*R* (*R* indicates the radial coordinate in the reference configuration), *λ*_*z*_ prescribed based on experimental observations, and *λ*_*r*_ = *R*/(*rλ*_*z*_) due to incompressibility.

For the active stress contribution in a blood vessel, a circumferential orientation for smooth muscle cells is generally assumed, yielding 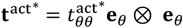. For a convenient energy function-type relation Ψ^act^ for the active contribution, we can also write

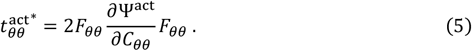

We then, for modelling convenience, split (Eq. 2) into passive Γ^pas^ and active Γ^act^ load bearing components due to the additive combination of the stress contributions according to

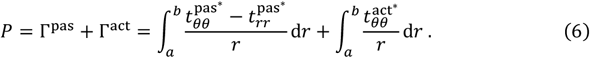

Eq. 6 allows us to evaluate the individual contribution of the passive and active stresses to the total transmural pressure. Since Ψ^pas^ is generally a hyperelastic constitutive relation that guarantees a monotonically increasing relationship between transmural pressure and inner radius for any given value of axial stretch, we focus on the active component of the pressure, analysing the potential of active stress models to lead to instability. Note that all constitutive models considered herein are phenomenological and do not consider the specific kinetics of calcium handling and actomyosin activity that allow for contraction.

### 2.1. Constant stress active contraction models

To capture experimental data on changes in vessel diameter with active contraction, we first demonstrate the approach with a simple one-parameter constitutive equation [5]. These models assume that the active contraction translates into a constant stress contribution, although this may be defined in different vessel’s configuration according to the specific model.

For a constant active stress in the current configuration (i.e., a constant active Cauchy stress), 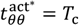 (with 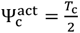 In 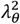), and we find

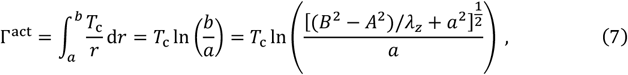

where *a* and *A* are the loaded and unloaded internal radii and *b* and *B* are the loaded and unloaded outer radii of the cylinder. Note that, for the incompressibility condition, 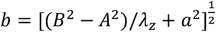. Furthermore, we assume the vessel axial stretch *λ*_*z*_ to be constant during the vessel pressurisation and, hence, independent from *a*. This assumption is consistent with the observation that most arteries (excluding the ascending aorta) undergo negligible axial deformations *in vivo* [19], as well as with commonly applied *ex vivo* testing protocols [5,17]. To evaluate whether this active stress formulation is prone to instability, we can then calculate the derivative of Eq. 7 with respect to the loaded internal radius of the cylinder *a* as

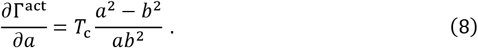

Given that 0 < *a* < b, 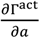 is negative for all deformations. Therefore, *P* = Γ^pas^ + Γ^act^ is a monotonically increasing function if and only if 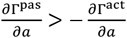 for all values of *a*. Although the occurrence of instability inherently depends on the functional form and parameter values of Ψ^pas^ as well as on *T*_c_, it is worth noting that the difference (*a*^2^ − *b*^2^) becomes more negative as *a* decreases and less negative as *a* increases.

Let us now consider an illustrative function of the form 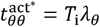 (with 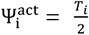 In *λ*_*θ*_), corresponding to a constant stress in the intermediate configuration (i.e., a constant 1^st^ Piola-Kirchhoff (PK) stress). Its associated active load bearing can be calculated as

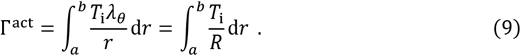

For the conservation of volume, *R* can be expressed as a function of the deformed radial coordinate *r*, so that the integral in Eq. 9 becomes

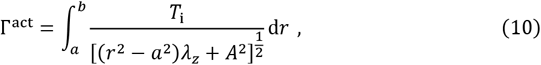

leading to the following relationship:

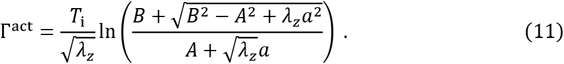

Eq. 11 allows then to evaluate changes in Γ^act^ with increasing luminal radius *a*

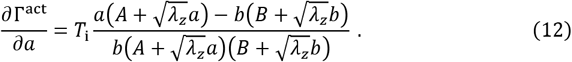

Once more, since 0 < *a* < *b* and 0 < *A* < *B*, the expression in Eq. 12 is negative for all values of *a* and *b*, and, hence, the constant 1^st^ PK active stress model is also prone to the development of instability.

Finally, let us consider a second illustrative case in which the smooth muscle contribution is modelled as a constant extra stress in the reference configuration (i.e., a constant 2^nd^ PK stress) [5], so that 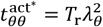 (with 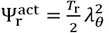). Once more, the active contribution to the intraluminal pressure is given by the integral

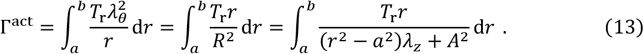

Integrating Eq. 13 yields

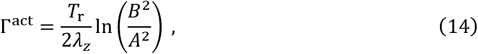

implying that the smooth muscle load bearing is independent from the deformed configuration (i.e., 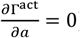). Therefore, provided that 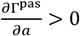 for all *a*, the present active stress model is not susceptible to instability issues.

### 2.2. Physiologically motivated models

The simplicity of constant active stress models makes them computationally convenient for application on sparse experimental smooth muscle contraction data and analytical analysis of stability characteristics. However, these models simplify the active mechanical contribution of vascular smooth muscle cells by neglecting their well-known parabolic force-length relationship [3,20]. For this reason, previous works have proposed more complex mathematical descriptions of active contraction, which account for the physiological mechanical behaviour of VSCMs.

In 1999, Rachev and colleagues [20] proposed the following expression to describe the VSMC stress contribution:

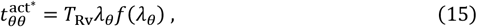

where, *T*_Rv_ is a VSMC stress-like parameter, *f*(*λ*_*θ*_) is a generic function capturing the physiological force-length relationship of the vascular smooth muscle, with max *f* = 1. Note that combining Eq. 15 and Eq. 5 yields the general expression for 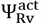:

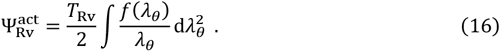

Different formulations of *f*(*λ*_*θ*_) have been proposed in the literature [20,21]. Here, we focus on a case which presents a convenient analytical solution.

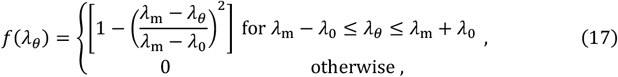

where *λ*_m_ indicates the circumferential stretch level for the maximal stress and *λ*_0_ controls the width of the bell-shaped function *f* (with *λ*_m_ ≠ *λ*_0_). Combining Eqs. 6, 15, and 17 and rearranging leads to

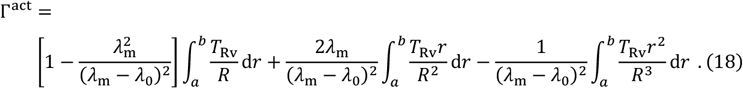

Integrating Eq. 18 (also see Eqs. 9 and 13) yields

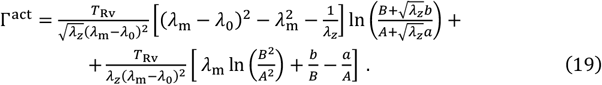

Differentiation with respect to *a* leads to

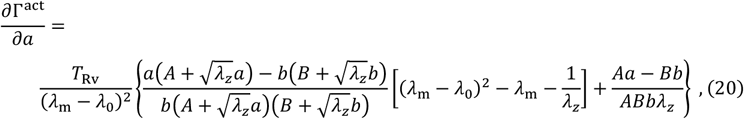

which is positive for

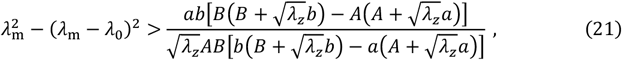

where all model parameters (i.e., *λ*_m_ and *λ*_0_) appear on the left-hand term and all geometrical features (i.e, *A* and *B*) and the deformation state (i.e., *a* and *λ*_*z*_) appear on the right-hand term. Since the left- and the right-hand terms are always positive, the sign of 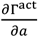 depends on the specific values of the active model parameters together with the deformation state and the unloaded vessel dimensions. Nonetheless, it is worth noting that the left-hand term of Eq. 21 increases with *λ*_0_ approaching *λ*_m_, while the right-hand term decreases with increasing *B*/*A* and *λ*_*z*_.

Zulliger et al. [22] proposed the following alternative active energy density function to capture the mechanical behaviour of the contracted VSMCs:

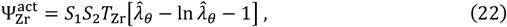

where 0 < *S*_1_ < 1 defines the degree of smooth muscle contraction (with *S*_1_ = 0 and *S*_1_ = 1 indicating the fully relaxed and maximally contracted states), *S*_2_ controls the deformation range in which VSMCs are able to generate force (with *S*_2_ = 1 for 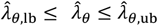 and 0 otherwise), *T*_Zr_ is a stress-like parameter and 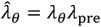 is the circumferential stretch acting on the VSMCs (with *λ*_pre_ defining their deposition stretch). Unlike Rachev’s model, this model captures only the ascending part of the VSMC force-length relationship, under the assumption that VSMC operates in this regime in the physiological pressure range. In this work, we focus on the behaviour of the arterial wall in the maximally contracted state (i.e., *S*_1_ = 1); i.e., when it is the most likely for modelled instable behaviours to occur. Differentiating Eq. 22 yields

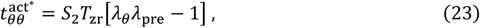

so that Eq. 23 reduces to the difference between the contributions of a constant 1^st^ Piola-Kirchhoff stress, *T*_*i*_ = *S*_2_ *T*_Zr_λ_pre_, and of a constant Cauchy stress, 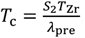. Using Eqs. 8 and 12, we find

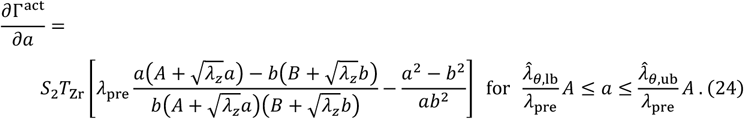

It follows that, in this deformation range, 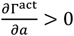 is guaranteed for

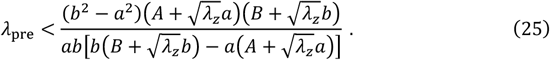

It can be shown that the term on the right-hand side of Eq. 25 monotonically decreases with increasing *a* (See thin black line in Figure 4, Panel E).

Finally, we consider the active energy density function recently proposed by Franchini et al. [23].

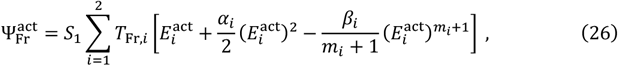

where

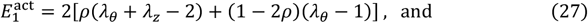

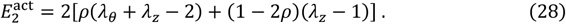

Eq. 26 accounts for the superimposed contribution of two families of vascular smooth muscle cells with a preferential circumferential and axial orientation, respectively. *S*_1_ ∈ [0,1] controls the level of smooth muscle activation, *T*_Fr,1_ and *T*_Fr,2_ are stress-like parameters for the circumferential and axial VSMC families, respectively. *α*_*i*_, *β*_*i*_, and *m*_*i*_ (with *i* = 1,2) control the shape of the force-length relationship of VSMCs; *α*_*i*_ controls the initial slow growth of active stress with increasing deformation and *β*_*i*_ and the integer *m*_*i*_ determine the rate of decrease in active stress at high deformations. *ρ* controls the VSMC orientation dispersion in the circumferential-axial plane (with *ρ* = 0 and *ρ* = 1/2 indicating no dispersion and full dispersion, respectively). Compared to the models by Rachev and Hayashi [20] and Zulliger et al. [22], Franchini’s model allows for more complex, non-symmetric shapes of the force-length relationship of VSMCs.

Following from our assumption that the VSMC contribution is purely circumferential to compare behaviour across models, we restrict our analysis to the case in which *T*_Fr,2_ = 0 and *ρ* = 0. Furthermore, by setting *S*_1_ = 1, we investigate maximal contraction. This leads to

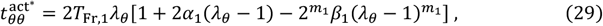

which, using Eqs. 11 and 14, becomes

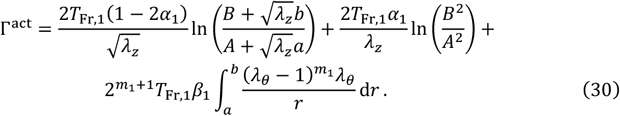

The analytical solution of Eq. 30 inherently depends on the value of the exponent *m*_1_. Here, we analyse the case *m*_1_ = 2 which presents a tractable analytical solution:

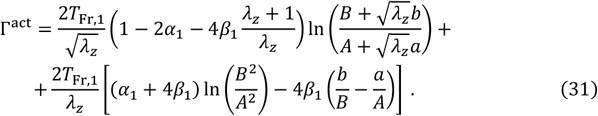

Differentiating Eq. 31 then leads to

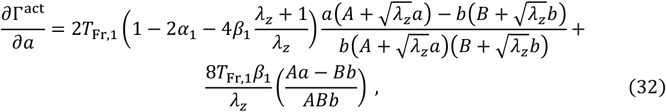

which is positive for

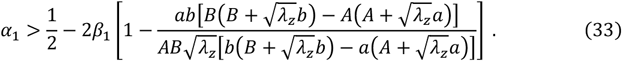

Note that the right-hand term in Eq. 33 grows with increasing *a* (thin black line in **Figure 4**, Panel F) and *λ*_*z*_.

## 3. Computational simulations

### 3.1. Mouse thoracic aorta

#### 3.1.2. Experimental data and parameter estimation

The overall aim of these computational simulations is to present case studies in which the aforementioned active stress models are used to capture experimentally measured contraction data. As the first case study, we chose representative data on the phenylephrine-induced contraction of the thoracic aorta of a wild type mouse, taken from a previous publication (Figure 2) [5]. In that work, segments of the mouse descending thoracic aorta were subjected to a comprehensive biaxial mechanical characterisation in passive conditions (i.e., with fully relaxed VSMCs). Then, while keeping the artery inflated at constant intraluminal pressure of 90 mmHg and axially stretched to its *in vivo* axial length, a bolus of 1 μM of phenylephrine was added to the organ bath to induce VSMC contraction. The induced reduction of the outer diameter was tracked over a period of 15 minutes (Figure 2).

**Fig. 2.**
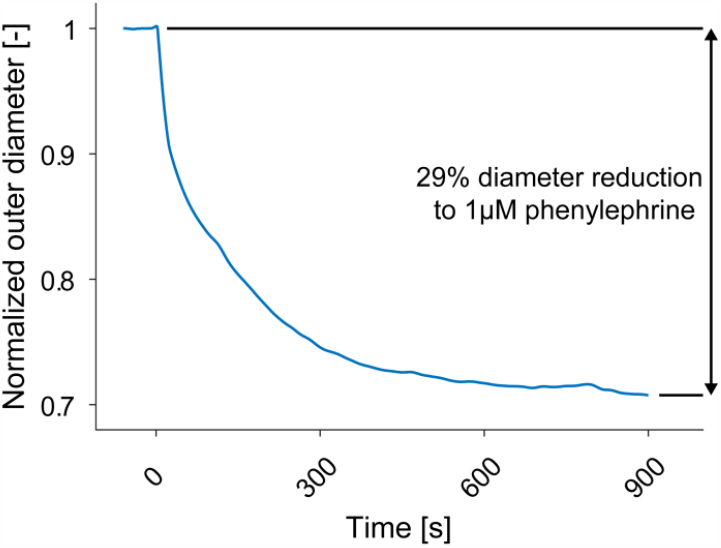
Experimentally measured contraction of the descending thoracic aorta of a male wild-type (C57BL/6J) mouse following addition of 1 μM of phenylephrine to the vessel bath. Throughout the 15 min-long experiment, the artery was axially stretched to its in vivo length and pressurised at 90 mmHg. Data from Spronck et al. [5].

To simulate this contraction, the arterial wall was modelled as a constrained mixture of elastin, VSMCs and four families of collagen fibres. A four-fibre family strain energy density function was used to capture the passive (i.e., elastin + collagen) mechanical behaviour of the mouse descending thoracic aorta.

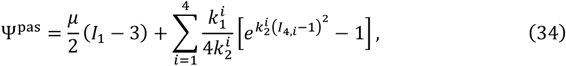

where *μ* is an elastin stiffness-like parameter, *I*_1_ = tr(**C**) is the first invariant of the right Cauchy-Green tensor, and 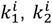 and 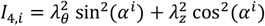 are a stiffness-like parameter, a non-linearity parameter the fourth-invariant of the right Cauchy-Green tensor for the *i*^th^ family of collagen fibres, with *α*^*i*^ = 0°, 90° and ±*α* for axially, circumferentially and diagonally oriented fibres, respectively. The parameter values of Ψ^pas^ (reported in Supplementary Information, **Table S1**) were obtained by averaging the modelled behaviour of mice in the C57BL/6J control group in our previous work [5] and refitting the parameters of Ψ^pas^ to the averaged curves.

The six active contraction models discussed above were used to capture the 29% phenylephrine-induced reduction in outer diameter. It is worth noting that, as shown in **Figure 3**, the results of this contraction experiment provide only a single mapping point between the mechanical behaviour in the fully relaxed and fully contracted states (i.e., at *in vivo* axial length and 90 mmHg of intraluminal pressure). It follows that a unique parameter solution can be identified only for those active stress models that encompass a single model parameter, namely the constant Cauchy, 1^st^ PK and 2^nd^ PK stress models. For the remaining three active stress models, we opted for imposing all model parameters, except for the VSMC stress-like parameter *T*_*j*_ with *j* = {Rv, Zr, Fr}, to either values reported in the literature or values that would result in a physiologically meaningful scenario. For the Rachev and Hayashi model, we set *λ*_m_ = 1.60 so that VSMCs provide their maximum stress at physiological pressures (i.e., ∼120 mmHg) [3,24]. Furthermore, by imposing *λ*_0_ = 0.80 we ensured continuity of the first derivative of pressure in the pressure range 0–200 mmHg (i.e., ensuring that the active stress is not null in this range). For the Zulliger et al. model, *λ*_pre_ was set to 1.83 as previously reported for the aorta of a normotensive rat [22]. As for *λ*_0_, 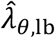 and 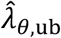 were assigned values that guarantee continuity of the first derivative of pressure in the range 0–200 mmHg (**Table 1**). For the Franchini et al. model, we chose *α*_1_ = 1.90 and *β*_1_=1.05, to yield, once more, the peak active stress at ∼120 mmHg [3,24]. Finally, the VSMC stress-like parameter (*T*_*i*_) of each model, with *i* = {c, i, r, Rv, Zr, Fr}, was estimated iteratively by minimising the error between the measured and modelled phenylephrine-induced reduction in vessel outer diameter (Π = *b*_measured_ − *b*_modeIIed_) using the MATLAB *lsqnonlin* function (MATLAB 2023a, MathWorks, Natick, MA, United States).

**Table 1.** Model parameters of the six active stress formulations used in study to capture the contractile behaviour of the mouse descending thoracic aorta.

| Model | Imposed parameters | Estimated parameters | Instability |
| --- | --- | --- | --- |
| Constant Cauchy | - | $T_c = 47.3 \text{ kPa}^*$ | Yes |
| Constant 1<sup>st</sup> PK | - | $T_i = 42.1 \text{ kPa}$ | No |
| Constant 2<sup>nd</sup> PK | - | $T_r = 37.8 \text{ kPa}$ | No |
| Rachev et al. [19] | $\lambda_m = 1.60$<br>$\lambda_0 = 0.80$ | $T_{RV} = 67.2 \text{ kPa}$ | No |
| Zulliger et al. [21] | $\lambda_{pre} = 1.83$<br>$\lambda_{lb} = 0.37$<br>$\lambda_{ub} = 4.21$ | $T_{Zr} = 45.3 \text{ kPa}$ | No |
| Franchini et al. [22] | $\alpha_1 = 1.90$<br>$\beta = 1.05$ | $T_{Fr} = 15.4 \text{ kPa}$ | No |

\*Note that the measured contraction data point falls within the instability region of the constant Cauchy active stress model (**Figure 3A**). Therefore, no value of $T_c$ is able to capture well the experimental data. The value of $T_c$ reported in this table corresponds to the value used in the simulation in **Figure 3A**.

**Fig. 3.**
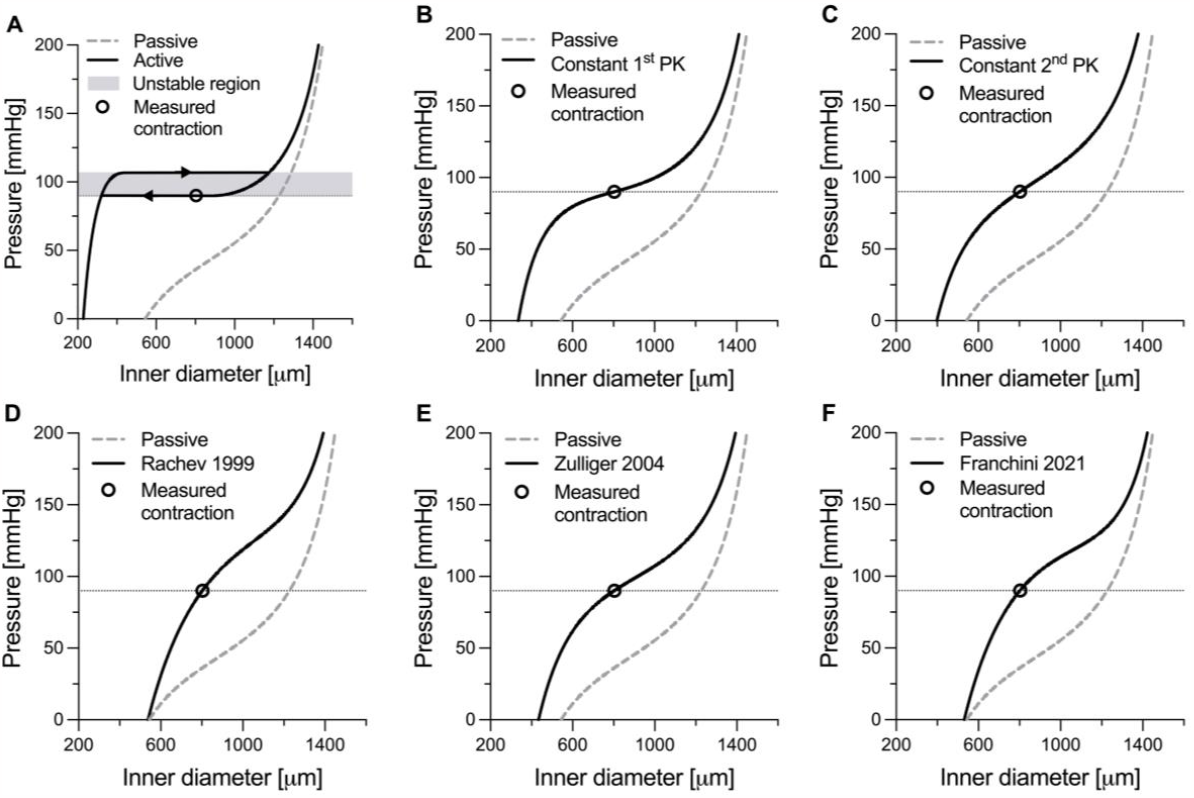
Simulated response to inflation at a constant (*in vivo*) value of axial stretch of the mouse descending thoracic aorta in both fully relaxed and maximally contracted conditions. The active behaviour was modelled using six different active stress formulations, namely A) constant Cauchy stress, B) constant 1^st^ Piola-Kirchhoff (PK) stress, C) constant 2^nd^ PK stress, D) Rachev model, E) Zulliger model, and F) Franchini model. The dashed grey lines and black circles in panel A–F indicate the passive behaviour of the wall (i.e., fully relaxed state) and measured contraction at 90 mmHg, respectively. The shaded area in Panel A indicates the region of instability. Dotted lines indicate 90 mmHg.

**Fig. 4.**
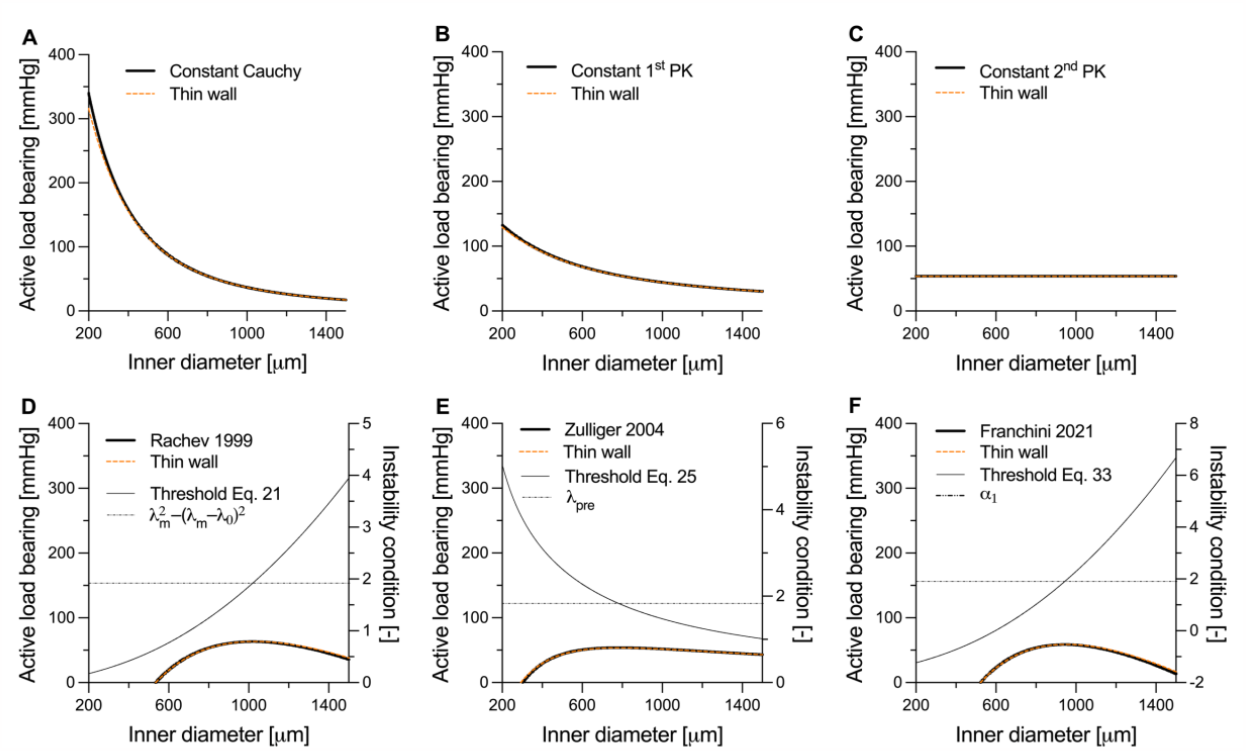
Active load bearing (Γ^act^) for the six active model considered in this study: A) constant Cauchy stress, B) constant 1^st^ Piola-Kirchhoff (PK) stress, C) constant 2^nd^ PK stress, D) Rachev model, E) Zulliger model, and F) Franchini model. Black solid and dotted-dashed lines in panels D–F indicate the left-hand and right-hand side of Eqs. 21, 25, and 33, respectively. In each graph, the intersection point between solid and dotted-dashed lines indicates where the first derivative of the active pressure changes sign. The red dashed lines in panels A–F indicate the approximated thin-wall solution (see Supplementary Information, Section S1).

#### 3.1.2. Results

The imposed and estimated model parameters of the six active stress models are reported in **Table 1**. The simulated response to inflation at the constant *in vivo* axial stretch of 1.60 is shown in **Figure 3**. The constant Cauchy stress model was the only active stress formulation that resulted in instability for the examined case study. As shown in **Figure 3A**, the measured contraction data point falls within the instability region and no-value of *T*_c_ can capture well the experimental data.

Figure 4. presents the isolated active load bearing Γ^act^ (**Figure 3**) for the six considered models. Despite not showing instability in the total pressure (i.e., *P* = Γ^pas^ + Γ^act^), all models except for the constant 2^nd^ PK active stress showed a negative Γ^act^–diameter slope in at least part of the investigated range of deformations (Supplementary information, **Figure S1**). As shown analytically, the 2^nd^ PK active stress model yields a pressure contribution that is independent from the deformation. As such, the pressure-diameter relationship in the contracted states preserves the same shape, although shifted upward, of the passive relationship. In agreement with our analytical derivation, *∂*Γ^act^/*∂a* was always negative for both the constant Cauchy and 1^st^ PK active stress models, particularly at small diameters (Supplementary information, **Figure S1A** and **B**). Given the high compliance of the fully relaxed aorta at small diameters, instability likely occurs in this diameter range when using these two models (e.g., **Figure 3A**).

The three physiology-driven models showed an opposite behaviour, with *∂*Γ^act^/*∂a* being negative only at high diameters (Supplementary information, **Figure S1D**–**F**). This is explained by the physiological bell-shaped VSMC force-length relationship that these models aim to replicate. Because Zulliger’s model captures only the ascending limb of this parabolic relationship, the resulting *∂*Γ^act^/*∂a* is less negative compared to both those of Rachev and Franchini (**Figure 4E** vs **Figure 4D** and **F** and Supplementary information, **Figure S1E vs Figure S1D** and **F**). Given the model parameters, vessel geometry and the deformation field, the deformation level at which *∂*Γ^act^/*∂a* changes sign can be found using Eqs. 21, 25, and 33 for Rachev’s, Zulliger’s, and Franchini’s model, respectively (intersection point between thin solid and dashed-dotted lines in **Figure 4D–F**). Nonetheless, because of the high passive stiffness of the vessel at high deformation levels, *∂*Γ^pas^/*∂a* > −*∂*Γ^act^/*∂a* in this range, so that no instability occurred in these simulations.

Figure 3. shows that all but the constant Cauchy model captured well the experimentally measured contraction of the mouse thoracic aorta at the luminal pressure of 90 mmHg. However, the absence of additional contractile data complicates evaluating whether the modelled contractile behaviour predicts well that of the artery for pressures below or above 90 mmHg. One aspect that may be considered is the modelled contraction at 0 pressure. VSMC’s contractile apparatus, particularly in large elastic arteries, is structured in such a way to provide its maximum force in the physiologically pressure range. As such, models that predict a very large reduction in diameter at 0 pressure (e.g., constant Cauchy and 1^st^ PK models, **Figure 3A** and **B**) are unlikely to closely match the actual contractile capacity of the vessel at low pressures. Nonetheless, these considerations are only qualitative in nature and more comprehensive experimental datasets, as the one proposed in the next example (see Section 3.2), are required to evaluate a model’s ability to capture the arterial contractile behaviour.

### 3.2. Contraction in dog arteries

#### 3.2.1. Experimental data and parameter estimation

In this second case study, we aimed to evaluate the ability of the aforementioned models to capture more extensive experimental data which encompass more than a single mapping point between the fully relaxed and maximally contracted state. In his seminal study on the contractile behaviour of large arteries, Cox [3] subjected segments of dog thoracic aortas and carotid, renal, iliac and mesenteric arteries to pseudo-physiological biaxial loading (pressurisation from 0 to 240 mmHg at the artery *in vivo* axial stretch). The biaxial experiments were first performed with the artery immersed in a physiological salt solution (PSS) and then repeated with a high potassium concentration PSS to measure the vessel’s mechanical behaviour in the fully relaxed and maximally contracted states, respectively. Sampling of the pressure-diameter relationships was performed at pressure intervals of 20 mmHg, with the diameter then reported as normalised with respect to the diameter at 0 pressure (*b*_0_) (Figure 3 in Cox [3]). To replicate these data, we first resampled both fully relaxed and maximally contracted pressure–normalised diameter relationships at the same normalised diameter values to yield the active load bearing 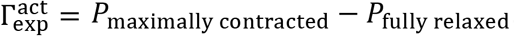. Because of the lack of detailed data on the tested vessels (e.g., unloaded thickness and diameter), we assume the vessel wall to be a thin membrane and approximate the average wall stretch (*λ*_*θ*_) with the normalised diameter (*b*/*b*_0_). Note that Supplementary Information, Section S1 illustrates how equivalents of the equations derived in Section 2 of this manuscript can be obtained for a thin-walled vessel. Furthermore, we assume values of radius-to-wall thickness ratios reported in the original manuscript (Table 2 in Cox [3]) to be representative of the vessel configuration at 0 pressure and the *in vivo* axial stretch. For each of the six active stress models presented above, the model parameters are then estimated by minimising the cost function

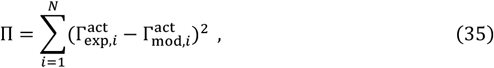

where *N* is the number of data points and the modelled active pressure is

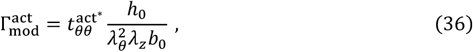

where *b*_0_/*h*_0_ is the radius-to-wall thickness ratio at 0 pressure, *λ*_*z*_ = 1 is the axial stretch (note that no further stretch is applied to the vessel during inflation), and 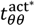 is the active Cauchy stress for the specific active stress formulation.

**Table 2.** Model parameters of the six active stress formulations considered in study to capture the contractile behaviour of five dog arteries.

| Model |  | Thoracic<br>aorta | Carotid<br>artery | Iliac artery | Mesenteric<br>artery |
| --- | --- | --- | --- | --- | --- |
| Constant Cauchy | $T_c$ [kPa] | 58.0 | 88.0 | 231.5 | 250.7 |
| Constant 1<sup>st</sup> PK | $T_i$ [kPa] | 48.2 | 75.9 | 195.8 | 216.0 |
| Constant 2<sup>nd</sup> PK | $T_r$ [kPa] | 36.0 | 60.3 | 155.8 | 177.6 |
| Rachev et al. [19] | $T_{Rv}$ [kPa] | 66.1 | 128.6 | 252.4 | 292.2 |
| | $\lambda_m$ [-] | 1.81 | 1.92 | 1.70 | 1.72 |
| | $\lambda_0$ [-] | 0.81 | 0.92 | 0.70 | 0.72 |
| Zulliger et al. [21] | $T_{Zr}$ [kPa] | 98.4 | 248.4 | 331.9 | 414.6 |
| | $\lambda_{pre}$ [-] | 1.25 | 1.09 | 1.40 | 1.35 |
| | $\lambda_{lb}$ [-]* | - | - | - | - |
| | $\lambda_{ub}$ [-]* | - | - | - | - |
| Franchini et al.<br>[22] | $T_{Fr}$ [kPa] | 13.0 | 14.9 | 69.1 | 74.0 |
| | $\alpha_1$ [-] | 1.4 | 2.5 | 0.9 | 1.1 |
| | $\beta_1$ [-] | 0.1 | 0.0 | 0.1 | 0.2 |
| | $m_1$ [-]** | 3 | - | 2 | 3 |

\* Missing values indicate that the estimated $\lambda_{lb}$ or $\lambda_{ub}$ exceeded the investigated range.
\*\* When $\beta_1 = 0$ the modelled behaviour is independent from $m_1$ . In this case, the value of $m_1$ cannot be estimated.

#### 3.2.2. Results

The experimental and modelled pressure-normalised diameter curves are presented in **Figure 5**, with the isolated active load bearing illustrated in **Figure 6**. The model parameters of the six active stress formulations for all the four arterial segments are reported in **Table 2**. As expected, for most arterial segments, the constant Cauchy and 1^st^ PK models poorly captured the experimental data, showing instability in the iliac (both), mesenteric (both) and carotid (only constant Cauchy) arteries. Despite its simplicity, the constant 2^nd^ PK model rendered a relatively good fit of the measured contractile behaviour for all arterial locations. Unsurprisingly, the fit of the experimental data further improved when the VSMC contraction was modelled using more complex physiology-driven models, which yield similar modelled behaviours independently from the chosen formulation.

**Fig. 5.**
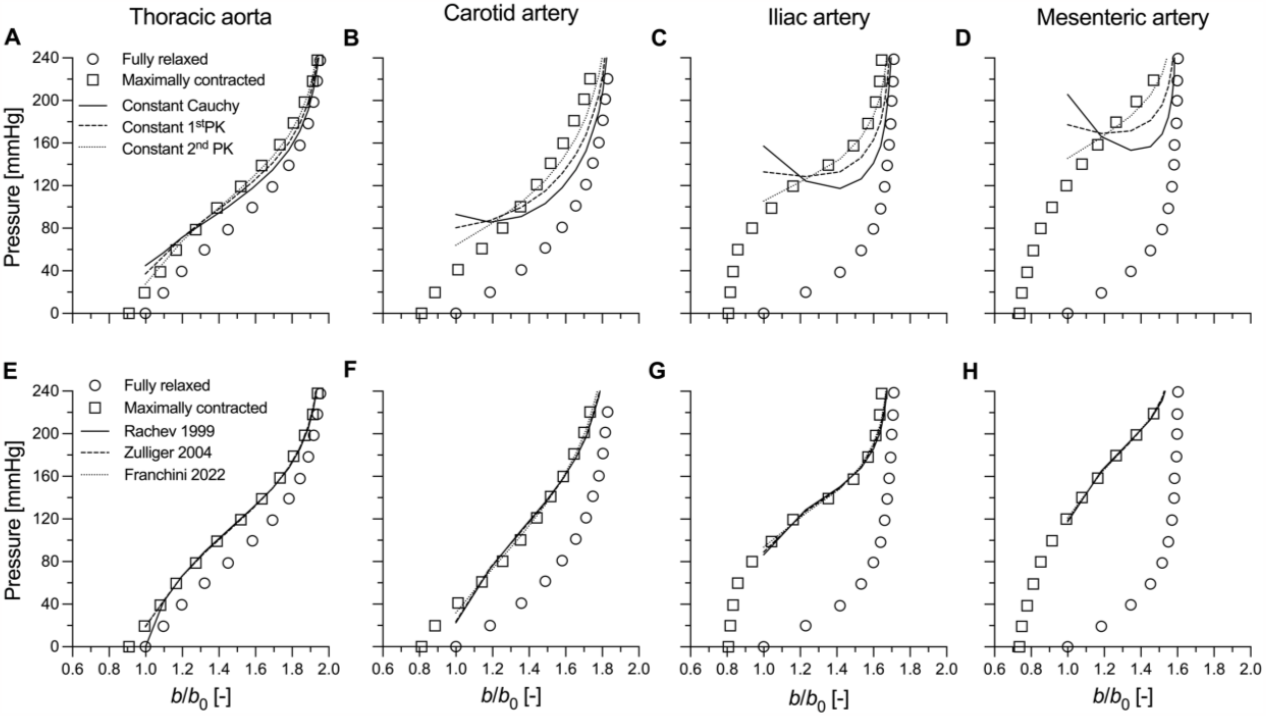
Experimental (adapted from Cox 1978) and simulated response to inflation at a constant (*in vivo*) value of axial stretch of the different canine arteries. Panels in the top row (A–E) present the fitted maximally contracted curves for the three constant stress models (i.e., constant Cauchy, 1^st^ Piola-Kirchhoff (PK) and 2^nd^ PK stresses). Panels in the bottom row (F–J) present the fitted maximally contracted curves for the three physiologically-driven models (i.e., Rachev and Hayashi [19], Zulliger et al. [21], and Franchini et al. [22]). *b*: loaded outer diameter; *b*_0_: outer diameter at pressure 0 mmHg.

**Fig. 6.**
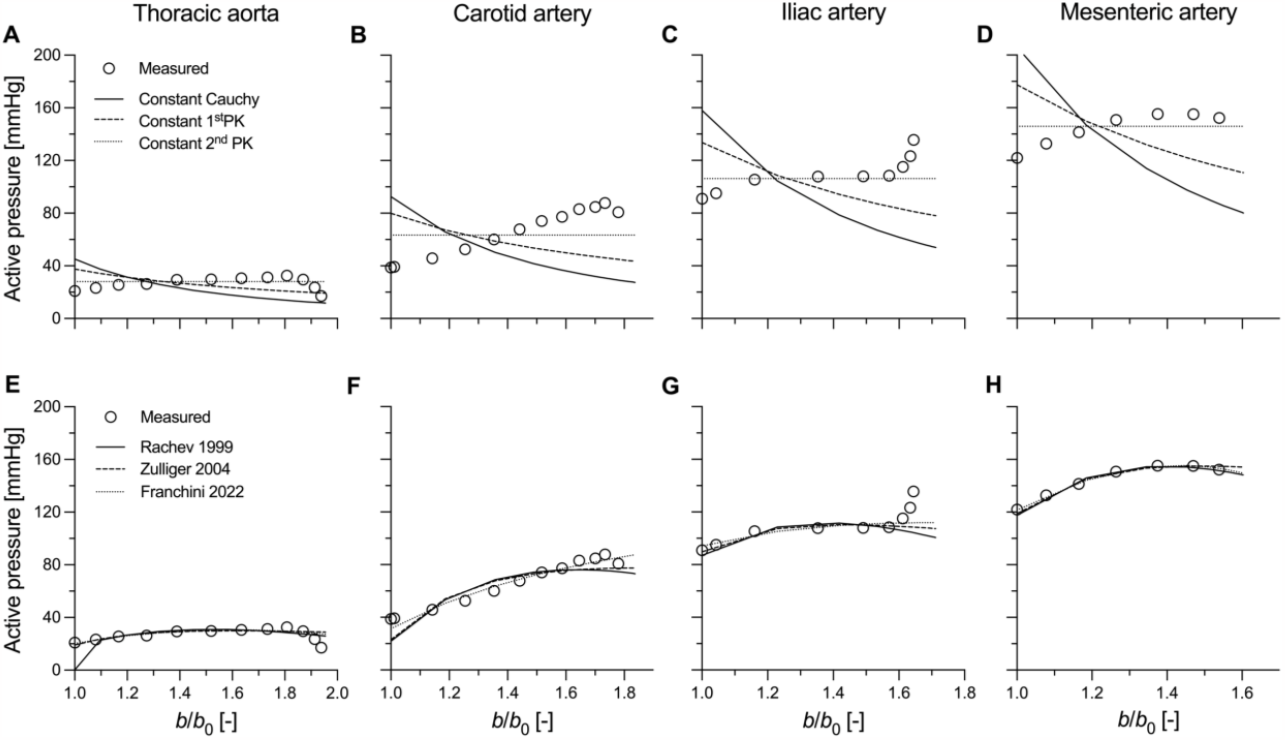
Experimental and simulated active pressure of the different canine arteries. Panels in the top row (A–E) present the fitted curves for the three constant stress models (i.e., constant Cauchy, 1^st^ PK and 2^nd^ PK stresses). Panels in the bottom row (F–J) present the fitted curves for the three physiologically-driven models (i.e., Rachev and Hayashi [19], Zulliger et al. [21], and Franchini et al. [22]). *b*: loaded outer diameter; *b*_0_: outer diameter at pressure 0 mmHg.

## 4. Discussion

Vascular smooth muscle cells play a key role in vascular homeostasis; their ability to modulate their phenotype and adapt their active tone during disease processes can both mitigate and contribute to pathological outcomes [1,5,13,14]. Therefore, devising effective methods to experimentally assess and then computationally capture their mechanical behaviour is key to predicting outcomes in adaptations. The experimental assessment of vasoreactivity is hindered by practical limitations, which yield paucity of data to inform computational models of active contraction. For this reason, in the present study, we analysed, both theoretically and computationally, six illustrative, phenomenological constitutive descriptions of active stress in blood vessels, evaluating their applicability to limited contraction data as well as their potential to model physiologically relevant behaviours.

Constant active stress models constitute the simplest approach to capture the mechanical behaviour of VSMCs. As they involve a single model parameter, these models can be applied to experimental data that include as little as a single active contraction data point [5]. The modelled response varies considerably depending on the configuration in which the constant stress is applied. For the constant Cauchy and 1^st^ PK active stress models, the VSMC contribution to the total pressure monotonically decreases with increasing deformation; notably *∂*Γ^act^/*∂a* is negative at small deformations. As relaxed arteries are highly compliant in this deformation range (i.e., *∂P*^pas^/*∂a* is small), these models may yield unstable contractile behaviours, especially when the measured contraction is strong (e.g., in peripheral muscular arteries). Indeed, in our simulations, both models led to unstable contractile behaviours for the dog iliac and mesenteric arteries (**Figure 5C** and **D**). Furthermore, the constant Cauchy stress model also yielded unstable solutions in central elastic arteries (i.e., the mouse thoracic aorta and the dog carotid artery). Although previous works have experimentally shown arterial instability in *ex vivo* experimental settings, with contracted vessels exhibiting sudden uncontrolled expansions for very small changes in transmural pressure [25–27], the average curves of the contracted dog arteries used herein did not show such behaviour. Hence, both constant Cauchy and 1^st^ PK stress models poorly captured the measured effect of VSMC contraction in dog arteries (**Figures 5A**–**E** and **6A**–**E**). Evaluating instability is harder for the mouse experimental data which included a single active contraction data point. Note, however, that this data point was experimentally measured at a near-steady state (**Figure 2**), which implies that it cannot be in an (experimental) unstable region. The constant Cauchy stress model thus seems less relevant also for the murine case study (**Figure 3A**).

Among all the considered models, the constant 2^nd^ PK active stress model represents a practically useful case with limited prior investigation. Beside implying a constant active stress contribution in the reference configuration, it also yields a constant active contribution to pressure (Eq. 14 and **Figure 4C**). Therefore, the modelled contracted behaviour preserves the shape of the passive behaviour and cannot result in instability. Despite its simplicity, this model captured the contractile behaviour of dog arteries relatively well (**Figures 5A**–**E**). Although the measured active contribution to pressure was parabolic in shape at most arterial locations, its variation over the investigated deformation range was relatively small. Hence, our simulations suggest the simplified constant 2^nd^ PK model may be advisable when capturing sparse contractile experimental data (e.g., murine thoracic aorta).

More complex multi-parameter models of active contraction aim to capture the physiological mechanical behaviour of VSMCs [20,22,23], which in turn is determined by the micromechanical interaction between actin and myosin filaments [28–30]. Indeed, contractile cells can generate their maximum force when subjected to a stretch that allows for the maximum number of cross-bridges between the actin and myosin filaments. A deviation in either direction from this optimal stretch level results in a progressively decreasing contractile capacity [28–30]. At the tissue level, this cell micromechanics results in the well-known parabolic force–length or pressure– diameter relationship of VSMCs (**Figure 4D**–**F** and **Figure 6F**–**J**). Overall, all three multi-parameter models of active contraction showed good ability to capture the contractile behaviour of all the canine arteries in the study of Cox [3]. As shown both theoretically and computationally, all three models have the potential to yield unstable modelled behaviours (i.e., *∂*Γ^act^/*∂a* < 0 in part of the deformation range), as could be expected by their reproduction of a parabolic force-length behaviour. Nonetheless, because the microstructural arrangement of VSMCs within the wall is such to guarantee peak active force/pressure generation around physiological/high physiological pressures [3], instability issues may only arise at deformation levels which are above this preferred VSMC working range. Note, however, that arteries exhibit high passive stiffness in this physiological deformation range, partially due to the recruitment of collagen fibres in the adventitia [5,31,32]. As such, the passive contributions make these contraction models robust against numerical/mathematical instability. Indeed, snap-through experimental behaviour indicative of a limit-point instability has been observed in muscular arteries when active stiffness is very high, but generally only with non-physiological pressurisation [28].

In the present study, we compared the performance of six mathematical descriptions of active VSMC contraction in blood vessels, with a particular interest towards their potential for mathematical instability. The three multi-parameter models that reproduce the physiological parabolic mechanical contribution of VSMCs showed good ability to capture experimental contractile data, as well as yield stable modelled behaviours in the physiologic loading range. Their use is, hence, recommended whenever an extensive experimental characterisation of the contractile behaviour of the studied vessel is possible. When such an extensive experimental characterization is not possible, our simulations show that the constant 2^nd^ PK active stress model provides a good balance between model accuracy and applicability, also preventing the risk of incurring mathematical instability.

## 5. Acknowledgements

## 6. Declarations

### Competing interests

The authors have no relevant financial or non-financial interests to disclose.

## Supplementary Information

### S1. Theoretical analysis in a thin-walled cylinder

For a thin-walled cylindrical tube (i.e., *b* − *a* ≪ *a*), Eq. 2 can be approximated to

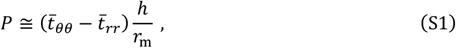

where *h* = *b* − *a* and *r*_m_ = (*a* + *b*)/2 are the wall thickness and the mid-wall radius, and 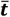 is the average Cauchy wall stress tensor across the wall thickness. Note that, under the assumption that 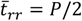, Eq. S1 is equivalent to the well-known Laplace equation

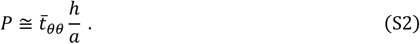

As done for the thick-wall derivation, the Cauchy stress in Eq. S2 can be split into an active and a passive contribution, to yield

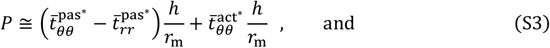

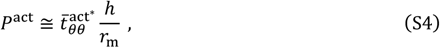

#### S1.1. Constant stress active contraction models

We then proceed with analysing the thin-wall equivalent of the three constant active stress models described above for the thick-wall case. For a constant active stress in the current configuration, 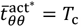, so that

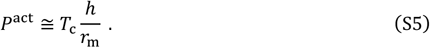

Knowing that, if *b* − *a* ≪ *b* + *a*, ln(*b*/*a*) ≅ 2(*b* − *a*)/(*b* + *a*), it becomes apparent that Eq. 7 can be recovered from Eq. S5 and that the constant Cauchy active stress model preserves the features discussed for the thick-walled case.

Similarly, for a constant active stress in the intermediate configuration, 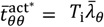 and

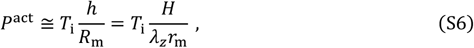

with *H* = *B* − *A* and 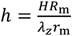 due incompressibility. Multiplying and dividing Eq. S6 by 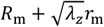 we obtain

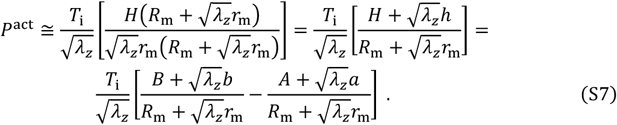

Once more, if, as in a thin-walled cylinder, 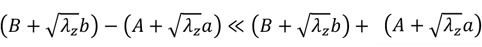, Eq. S7 is equivalent to Eq. 11.

Finally, for a constant active stress in the reference configuration, 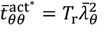, leading to

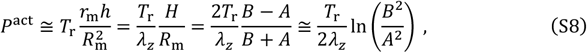

With In 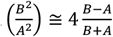 if *B* − *A* ≪ *B* + *A*.

#### S.1.2. Physiologically motivated models

With Eqs. S5, S6, and S8, we have shown that thin-walled modelling does not alter the behaviour of the considered constant active stress models, including the potential development of instability. As more complex models are typically equivalent to the superimposed contribution of these simpler formulations, the same argument holds also in those more complex cases. For a thin-walled vessel, the Rachev and Hayashi’s active component of the intraluminal pressure is

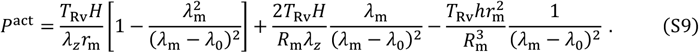

Recalling Eq. S7 and using the incompressibility constraint, Eq. S11 can be rewritten as

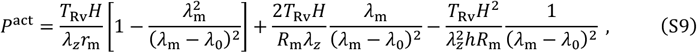

Noting that in a thin-wall cylinder *H*^2^/*hR*_m_ ≅ 0 ≅ *H*/*r*_m_, Eq. S12 can be simplified to

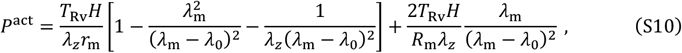

which, considering 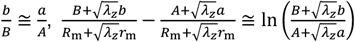, and 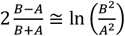, is equivalent to Eq. 19.

As seen in Eq. 23, the Zulliger et al. active stress model reduces to the subtraction between a constant 1^st^ Piola-Kirchhoff and a constant Cauchy stress term. In the thin-walled form, the active pressure is

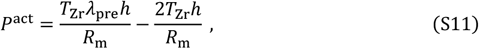

which recalling Eq. S7 and S8, for a thin-walled cylinder approximates well

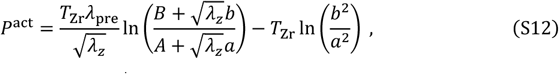

i.e., the thick wall expression of *P*^act^.

The thin-walled expression of the Franchini et al.’s model active pressure is

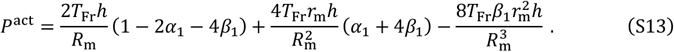

Recalling Eq. S7 and using the incompressibility constraint, Eq. S13 can be rewritten as

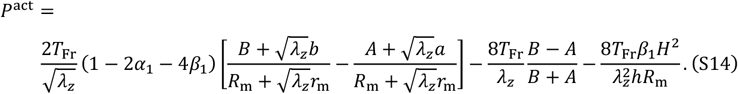

As for Eq. S9, in a thin-walled cylinder, Eq. S14 can be simplified to

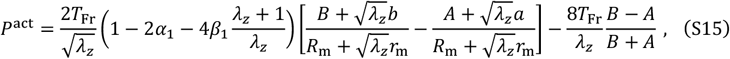

which, considering once more 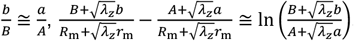, and 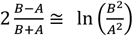, is equivalent to Eq. 31.

## Supplementary tables

**Table S1.** Four-fibre family strain energy density function parameters (taken from averages of the C57BL/6J control group in Spronck et al. [5]).

| $\mu$[kPa] | $k_1^1$ [kPa] | $k_2^1$ [-] | $k_1^2$ [kPa] | $k_2^2$ [-] | $k_1^{3,4}$ [kPa] | $k_2^{3,4}$ [-] | $\alpha^{3,4}$ [°] |
| --- | --- | --- | --- | --- | --- | --- | --- |
| 26.9 | 4.5 | 0.36 | 11.6 | 0.05 | 0.9 | 1.04 | $\pm 33$ |

## Supplementary figures

**Figure S1.**
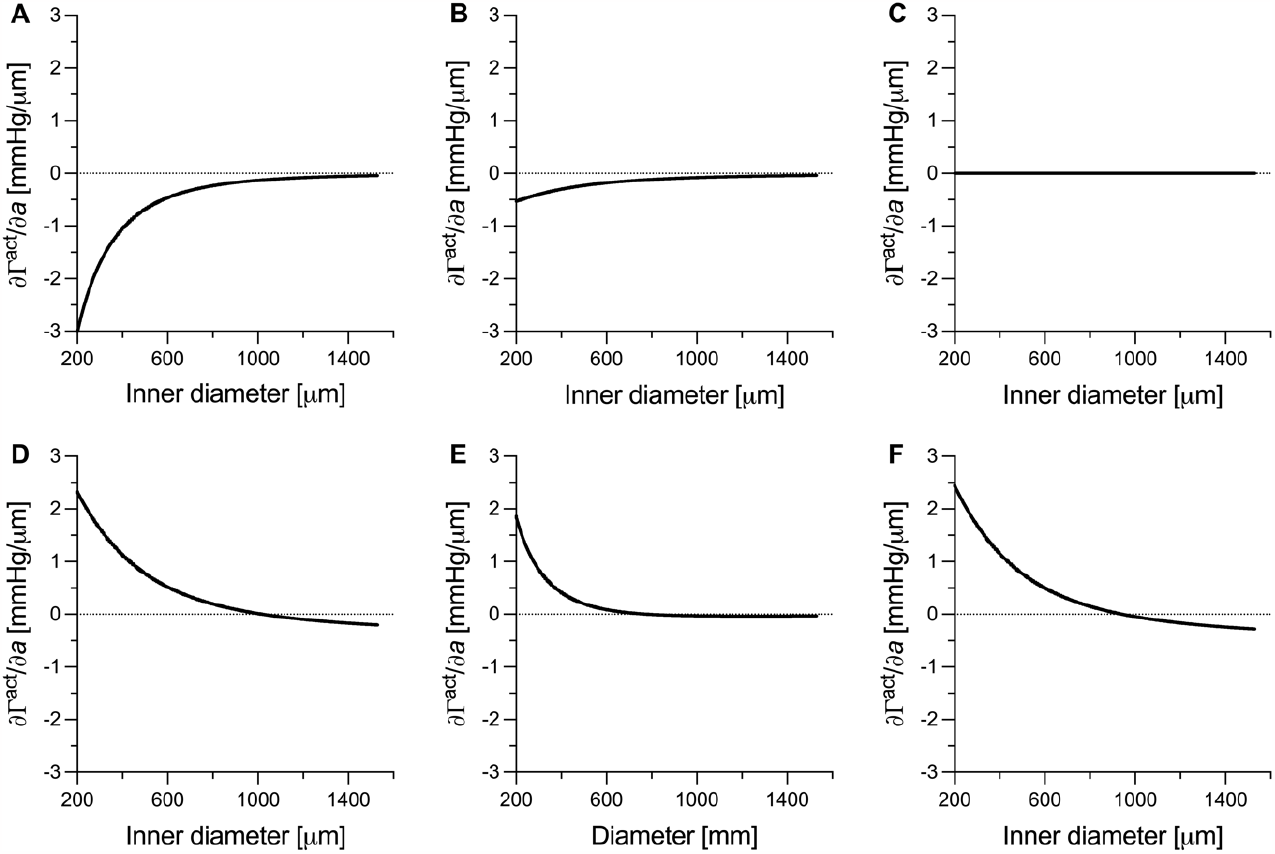
Slope of the active load bearing (Γ^act^) – inner radius (*a*) relationship of the six active stress model used to capture the contractile behaviour of the mouse thoracic aorta: constant Cauchy model (Panel A), costant 1^st^ Piola Kirchhoff (PK) model (Panel B), constant 2^nd^ PK model (Panel C), Rachev model (Panel D), Zulliger model (Panel E), and Franchini model (Panel F). The modelled contractile behaviour may show instability whenever the slope *∂*Γ^act^/*∂a* is negative, depending on the slope of the passive load bearing (Γ^pas^) – inner radius (*a*) relationship.

## Notes

### Competing Interest Statement

The authors have declared no competing interest.

